# Particle size determines mucociliary transport mechanisms in normal and cystic fibrosis airways

**DOI:** 10.64898/2026.07.01.735890

**Authors:** Michael Scott, Kaleb C. Bierstedt, Weijie Du, Mitchell J. Riley, Anthony J. Fischer, Yuliang Xie

## Abstract

A wide spectrum of microparticles is inhaled with each breath, deposited on airway surfaces, entrapped in the mucus, and removed by mucociliary transport (MCT). However, the influence of particle size on MCT remains largely unknown. Here, we investigated the MCT of microparticles with a trachea-on-a-chip method that integrates a micro-machined device with a trachea explant from newborn pigs. This method preserves airway structures for mucus secretion and cilia beating (*e.g.,* airway surface epithelia and submucosal glands), maintains physiological air-liquid-interface on the airway surface, and allows tracks motion of microparticles with high resolution. Using this method, we found that, in normal airways, 6 µm polystyrene particles clear rapidly, whereas 102 µm particles clear slower and require mucus strands for motion. In cystic fibrosis (CF) airways, MCT of microparticles reduces, but particle size-dependence persists. Methacholine increases particle motion in normal airways, but not in CF airways. These findings suggest two distinct MCT processes, in which large particles rely on mucus strands for clearance, small particles can be cleared independent of mucus strands, and CF disrupts both.

Graphical Abstract

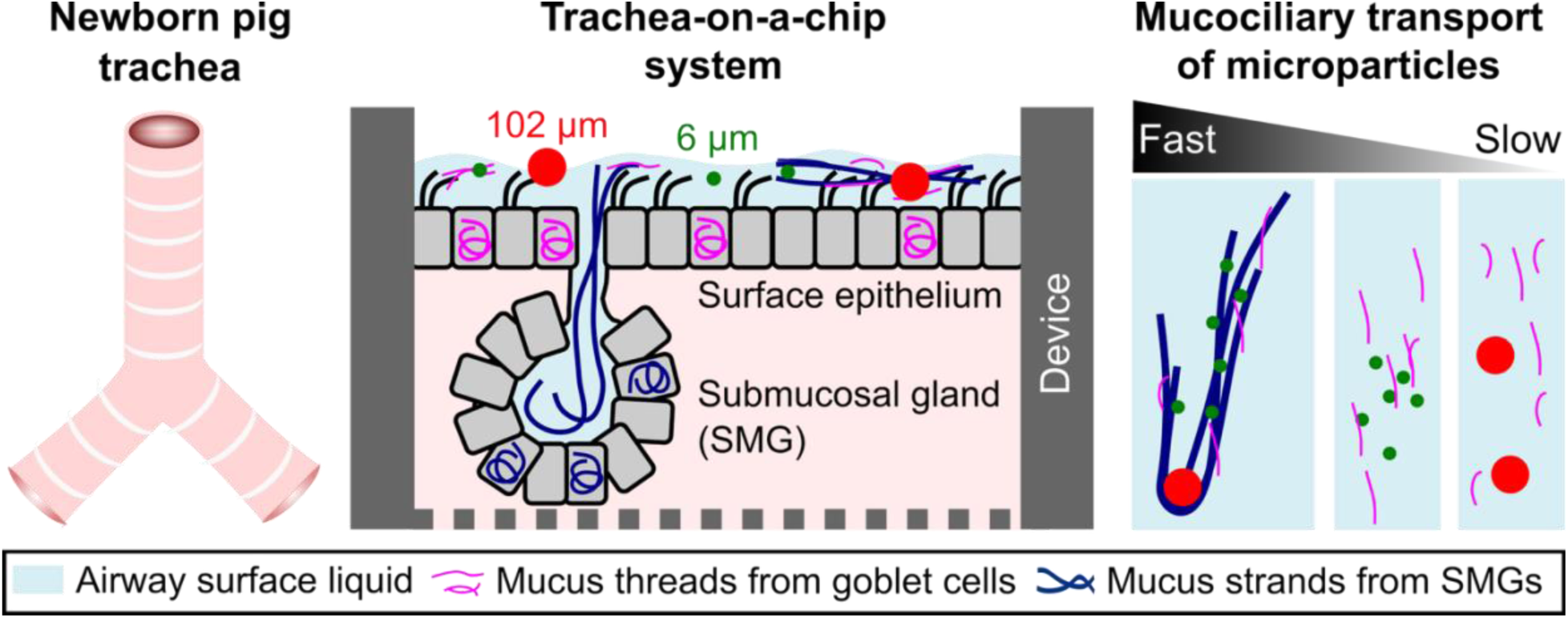

## Introduction

With each breath, a wide spectrum of microparticles (1-1000 μm in diameter) is inhaled, including dust (10-100 µm), particulate matter (PM_10_), and bacteria (1-10 µm). Due to their size, inhaled microparticles predominantly deposit in the conducting airways [1, 2], which include cartilaginous airways extending from the trachea for ∼16 generations to the respiratory bronchioles. The removal of inhaled microparticles from the conducting airways is through mucociliary transport (MCT) [3]. MCT is a process that traps microparticles in mucus secreted from airway surface goblet cells and submucosal glands (SMG) [4], and propels them out of the airways with synchronized ciliary beating. In airway diseases such as cystic fibrosis (CF) and primary ciliary dyskinesia, abnormal mucus secretion or ciliary beating impairs MCT [5–7], resulting in prolonged residence of microparticles/bacteria in the airways, exacerbating the risk of infection and inflammation [8]. Given the prevalence of inhaled microparticles and the role of MCT in determining their downstream impacts, it is important to understand how particle size influences MCT.

Insight into size-dependent MCT of microparticles would also have implications for the diagnostics and therapeutics of airway diseases. For example, various micro-radiotracers are used clinically to assess airway clearance functions [9], and nebulized aerosols and microparticles are inhaled to deliver medications to the airways [10, 11]. Knowledge of the size-dependence of MCT could improve the interpretation of diagnostic results and aid in developing effective drug carriers for airway diseases. However, previous studies on MCT of microparticles have been confounded by several factors. Clinical studies use the residence time of radiotracers to assess MCT, which is influenced by both their deposition positions and their motion on airways [7]. *In vivo* animal studies tracked the motion of large Teflon [12] and tantalum particles [13] (200-300 µm) on airway surfaces; however, tracking smaller particles remains challenging due to limitations in spatial and temporal resolution. *In vitro* methods depict particle motion on airway epithelial cell cultures [14]; however, cell cultures lack the complex cell types and tissue architecture, such as SMGs, that are key to MCT. To maintain airway structure and improve resolution, the MCT of microparticles has been investigated on tracheal explants from chicken embryos [15, 16] and pigs [17]. Nonetheless, most studies used submerged airway explants, where excessive liquid coverage destroyed the physiological airway mucus, thereby impacting the MCT of particles. As a result, the influence of microparticle size on MCT on airway surfaces with normal mucus remains largely uncertain.

We developed a “trachea-on-a-chip” system to investigate MCT of microparticles under normal and diseased airway conditions. The trachea-on-a-chip device integrates micromachined devices with a piece of trachea explant from newborn pigs that closely resembles human airways [18], including the presence of SMGs for mucus production and MCT [19]. It maintains the airway tissue in a non-submerged, air-exposed environment, thereby preserving a physiological mucus layer that facilitates MCT. It also enables precise control of microparticle deposition and allows high-resolution tracking of their motion *via* MCT. With the trachea-on-a-chip method, we compared MCT of microparticles with 6 μm and 102 μm diameters, representing inhaled bacteria and larger irritants, respectively. Since airway insults can elicit a cholinergic reflex that can increase ciliary beat frequency [20] and SMG secretion [21], we tested the cholinergic agonist methacholine (Mch) on the MCT of microparticles. To understand how disease impacts particle MCT, we studied airway explants from newborn wild-type (WT) and CF pigs. CF pigs spontaneously develop human airway disease hallmarks soon after birth [18]. The use of newborn pigs also avoids secondary disease manifestations such as infection, inflammation, and airway remodeling seen in older subjects.

## Results

### MCT of microparticles was investigated with a trachea-on-a-chip device

The experimental timeline to study MCT of microparticles is described (Fig. 1A). Before the experiment, a trachea segment (∼1.5 cm in length) was removed from newborn pigs, opened ventrally with a longitudinal cut, and mounted on the trachea-chip device (Fig. 1B). The micro-machined trachea-chip device contains four parts: a bottom part with a chamber (8 mm in length and 5 mm in width) for basolateral perfusion, a metal mesh to support the trachea explant, a top part, and a cover that together form an apical chamber (8 mm in length and 5 mm in width) for air perfusion, particle application, and observation of particle motion. When a trachea explant was mounted with the device, the apical and basal components were separated for individual perfusion. The whole trachea-chip device was aligned *via* screws and anchored on top of an optical breadboard.

**Fig. 1.**
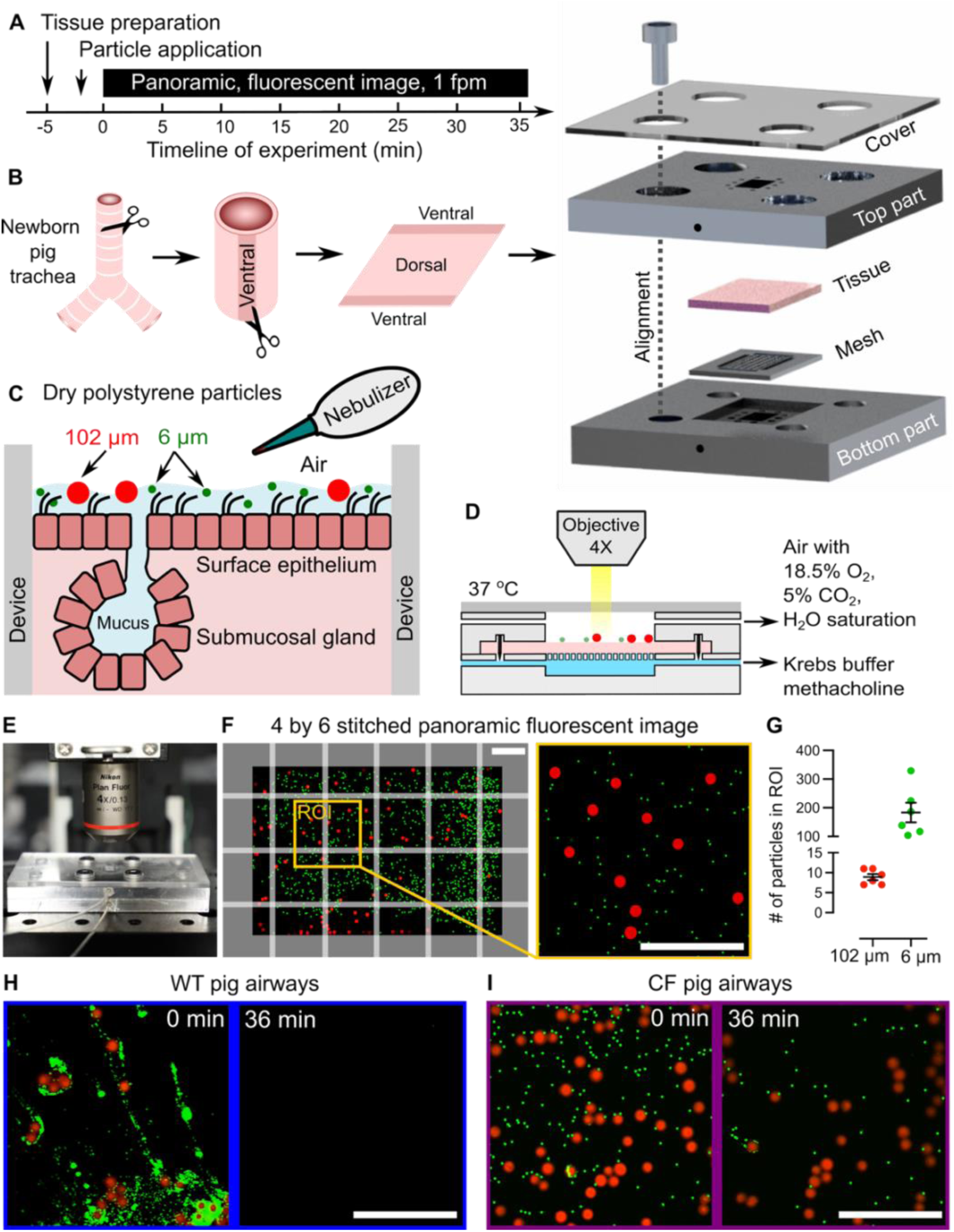
MCT of microparticles are investigated using a trachea-on-a-chip device. (A) Timeline of studying MCT of microparticles on a porcine trachea-on-a-chip device. (B) Schematic of assembling trachea tissue onto the trachea-on-a-chip device. (C) Dry polystyrene microparticles (6 μm diameter in green and 102 μm in red) were applied onto the airway surface using a homemade nebulizer. (D-E) The device was anchored in an environmental chamber under a confocal microscope. Air and medium perfusions were through the top and bottom chamber, respectively. (F) Panoramic, fluorescent images were used to visualize microparticles on the airway surface. To study particle motion, a region of interest (ROI, yellow box, 2 mm by 2 mm) was defined. (G) Number of microparticles applied within the ROI. (H-I) Representative images of microparticles at 0 and 36 min of the experiment on WT and CF airway surface. In panels F, H, and I, the scale bars are 1 mm. In panels F and I, green signals are digitally highlighted for visibility.

We applied dry fluorescent polystyrene microparticles (102 µm in red, 6 µm in green) onto the apical side of the airway using a homemade nebulizer (Fig. 1C and S1). After particle application, we perfused the apical chamber with ∼100% relative humidity air containing 5% CO_2_ to maintain a physiological mucus layer. We perfused the basolateral chamber with Krebs-Ringer buffer with HCO_3_ ^-^/CO_2_ to sustain tissue viability and function (Fig. 1D). In some experiments, the cholinergic agonist methacholine was perfused basolaterally to stimulate a cholinergic reflex [20]. The temperature of the airway explant, device, and perfusate was maintained at 37°C using an environmental chamber.

We used confocal microscopy to visualize the airway surface in the apical chamber (Fig. 1E), where panoramic images were stitched from 24 individual images (Fig. 1F). To analyze particle motion, a region of interest (ROI, 2 mm by 2 mm) was selected on the airway surface. The position of the ROI was chosen away from the edges of the apical chamber because sharp, demarcated edges impede the motion of particles nearby. Within the ROI, there were ∼10 of 102 µm particles, and ∼200 of 6 µm particles without significant particle aggregations (Fig. 1G). To study the motion of microparticles, panoramic images were captured every minute (*i.e.*, 1 fpm) for 36 min. In general, we observed that, by 36 min, particles were cleared entirely from the ROI on WT airways (Fig. 1H, Movie S1). At the same time point, many particles were retained on the airway surface ROI on CF airways (Fig. 1I, Movie S1), indicating defective MCT.

### MCT of microparticles was quantitatively assessed by particle clearance and speed

We investigated the motion of individual microparticles on the airway surface with the trachea-on-a-chip; Fig. 2A shows several example trajectories of 6 µm and 102 µm particles. We found that both 102 µm (red) and 6 µm (green) particles moved in the lung-larynx direction due to directional cilia beating (*e.g.*, upward direction, Fig. 2A), which is consistent with MCT *in vivo*. Using Imaris software, we conducted particle tracking, counting, and speed calculations every minute until all particles left the ROI or 36 minutes had elapsed (*i.e.*, the end of the experiment). In a representative particle tracking example on a WT pig airway, both 6 µm and 102 µm particles were successfully visualized (Fig. 2B), and a handful of trajectories are shown. Most particles were cleared from the ROI in ∼10 min, suggesting effective MCT in WT airways. Noticeably, 6 µm particles moved faster than 102 µm particles, evidenced by the red-shifted color of particle speed analysis along their trajectories (Fig. 2B).

**Fig. 2.**
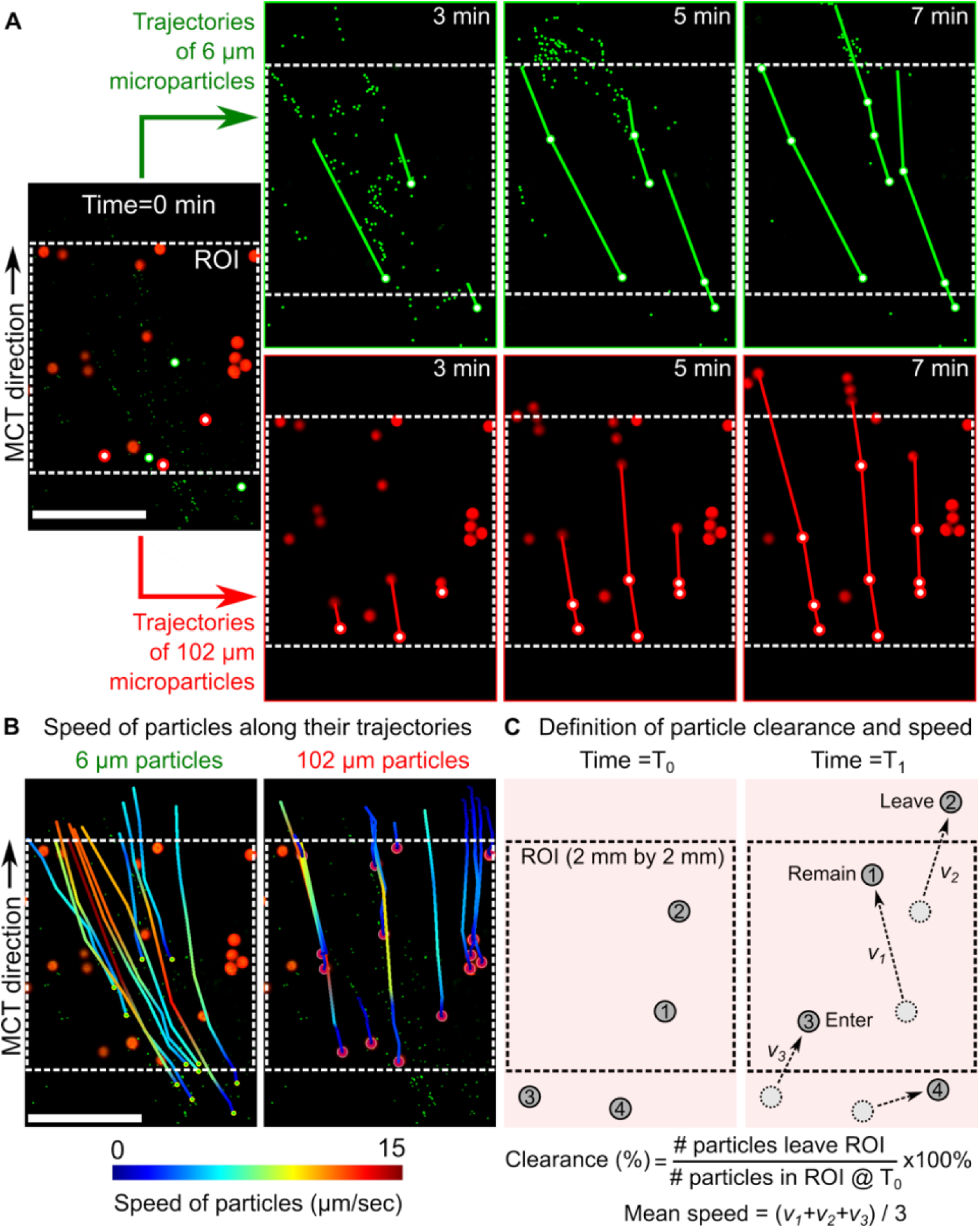
MCT of microparticles are visualized and quantified. (A) Image sequences showing the representative motions of 102 μm and 6 μm microparticles on a WT airway surface, where the region of interest (ROI, 2 mm by 2 mm) is defined to observe particle motion. (B) Computed particle trajectories and speeds with a color gradient to denote particle speed along each trajectory. 6 µm particles consistently moved faster than 102 µm particles. (C) Schematic demonstration of indicators for MCT of microparticles, where the particle clearance from the airways and the mean speed of particle motion were calculated for both microparticles. In panels A-B, the scale bars are 1 mm. In panel A, green signals are digitally highlighted for visibility.

We designed two indicators, *i.e.*, particle clearance and particle speed, to assess the MCT of microparticles (Fig. 2C). For example, at time T_0_, two particles (#1 and #2) are in the ROI, and two particles (#3 and #4) are outside the ROI. Between T_0_ and T_1_, particles move due to MCT; the displacement of each particle divided by the duration T_1_-T_0_ yields the speed of a particle.

Among particles that were tracked, we report the maximum speed (*v_1_*), median speed (*v_2_*), minimum speed (*v_3_*), and mean speed ([*v_1_*+ *v_2_*+*v_3_*]/3). At time T_1_, particle motion will generate one of four outcomes in relation to the ROI: particles that remain in (#1), leave (#2), enter (#3), and never enter (#4) the ROI. The particle clearance is defined as the percent of particles that leave the ROI between T_0_ and T_1_ (#2) relative to the number of particles in the ROI at T_0_ (#1 and #2). Of note, when calculating particle speed, particles that remain, leave, and enter the ROI are all considered. When calculating particle clearance, we only quantified particles that were initially within the ROI and ignored those that entered the ROI after T_0_. In experiments, the particle clearance and speed were calculated every 1 minute.

### Particle size, CF disease, and methacholine (Mch) stimulation impact particle clearance

We tested how 3 independent factors (*i.e.*, 6 µm vs. 102 µm particles, CF vs. WT, and Mch vs. basal) impact particle clearance and speed. These factors produced a total of 8 experimental groups. Each group contains 5 biological replicates. Each replicate produces the clearance and speed data from 30-50 particles on the airway surface.

We found that particle size impacts its clearance. On WT airways, 6 µm particles were cleared from the airway surface in ∼5 min under baseline conditions, and 102 µm particles had lower clearance than 6 µm particles (dot lines, Fig. 3A). To quantify how each factor impacts the particle clearance, we adopted the concept of hazard ratio using a Cox proportional hazard model and developed the indicator “clearance ratio”. For an individual factor, a clearance ratio >1 indicates that it increases particle clearance compared to its comparator, and vice versa. Using this method, we found that the clearance ratio of 6 µm vs. 102 µm particles was >1 in the strata of WT-basal, WT-Mch, CF-basal, CF-Mch, and all pooled data (Fig. 3C). This result confirmed that particles with a smaller size are cleared better on both WT and CF airways, with and without Mch stimulations.

**Fig. 3.**
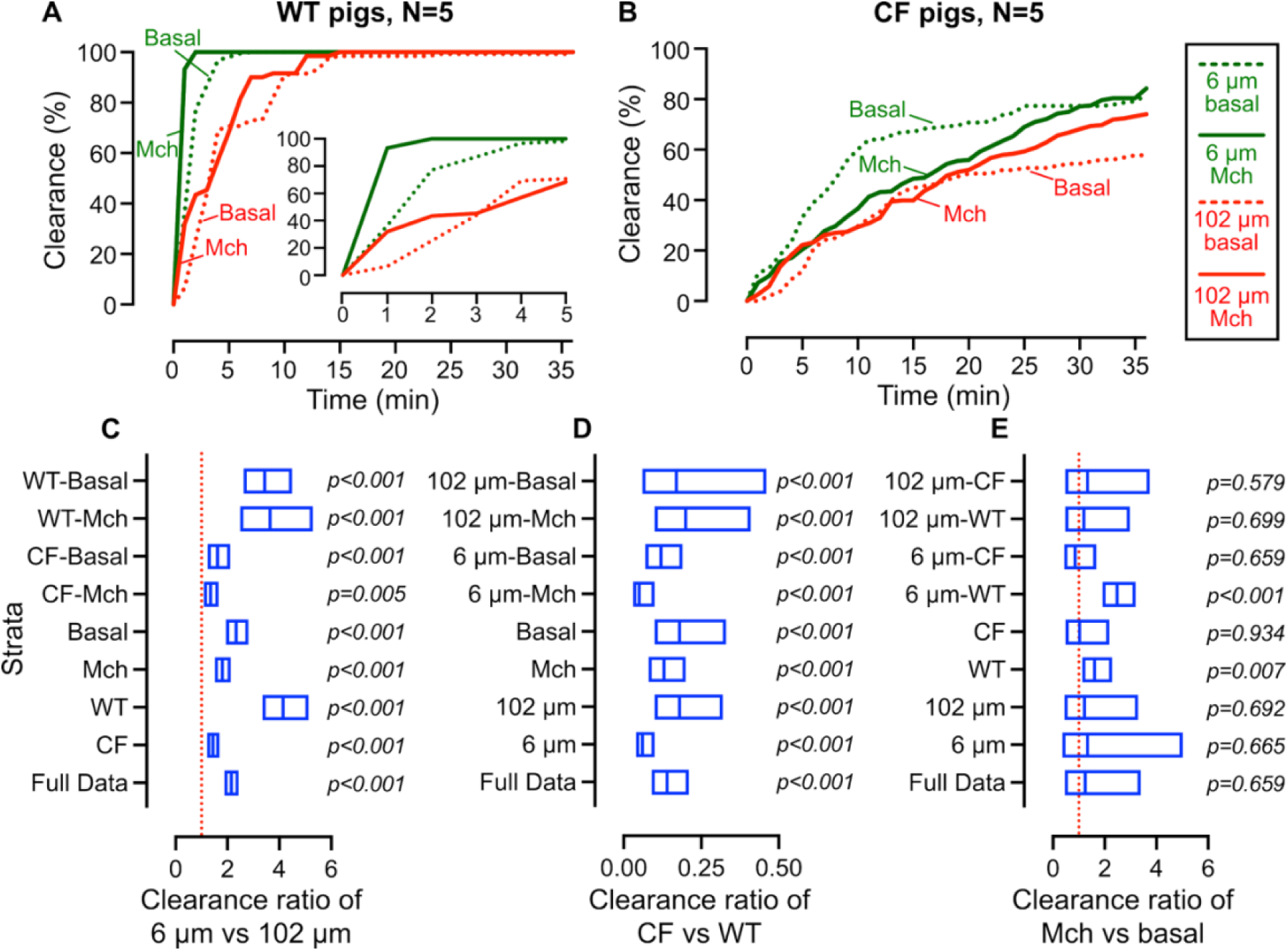
Particle size, CF airway disease, and methacholine (Mch) stimulation impact particle clearance from the airways. (A-B) Percent of particle clearance on wildtype (WT) and CF airways. (C-E) Effect of particle size, CF airway disease, and methacholine stimulation on particle clearance ratio. In panel A-B, each curve represents the mean value of particle clearance from 5 pig airways. In panel C-E, data are represented by clearance ratio, 95% confidence intervals, and p-values. Cox proportional hazards models were used to assess the effects of genotype (CF vs WT), particle size (6 µm vs 102 µm), and methacholine (Mch vs basal) on time of particle clearance on the airways. N=5 pigs for each condition.

Particle clearance on CF airways is lower than on WT airways, where a portion of particles remain on the airway surface by the end of 36 min (Fig. 3B). The effect of CF on particle clearance was quantified by the clearance ratio of CF vs WT airways. In all strata, including the full data set, the clearance ratio of CF vs WT was <1 (all p<0.001) (Fig. 3D), suggesting that CF caused defective microparticle clearance *via* MCT.

Cholinergic agonists Mch stimulate ciliary beating and SMG secretion; thus, we perfused it to the basolateral size of the tissue and tested its impact on particle clearance. We found that, on WT airways, Mch stimulation increases the clearance of 6 µm particles, but not 102 µm particles, compared to basal conditions (Fig. 3A). On CF airways, however, Mch seems ineffective on particle clearance (Fig. 3B). We quantified the effect of Mch by the clearance ratio of Mch vs basal conditions. For all WT airway data, the clearance ratio was >1 (p=0.007), suggesting Mch increases particle clearance via MCT (Fig. 3E). In contrast, for all CF airway data, the clearance ratio is ∼1 (p=0.934), suggesting that Mch’s effect on particle clearance is trivial. These results are consistent with our previous *in vivo* observations, where we found that Mch does not improve the clearance of tantalum micro-disks on CF pig airways [4].

### Particle size, CF disease, and Mch stimulation impact particle speed

Particle clearance is a consequence of their motion on the airways; thus, we analyzed the mean, maximum, median, and minimum speeds of particles. On WT airways, the mean speed of 102 µm particles was noticeably slower than that of 6 µm particles under basal conditions and with Mch stimulation (Fig. 4A). To quantify the speed difference, we calculated the mean speed ratio of 6 µm vs. 102 µm particles using a generalized linear mixed model. We found that the mean speed ratio of 6 µm vs 102 µm particles was >1 in all strata and pooled data (all p<0.05) (Fig. 4C). This result confirmed that smaller particles move faster in both WT and CF airways, with and without Mch stimulation.

**Fig. 4.**
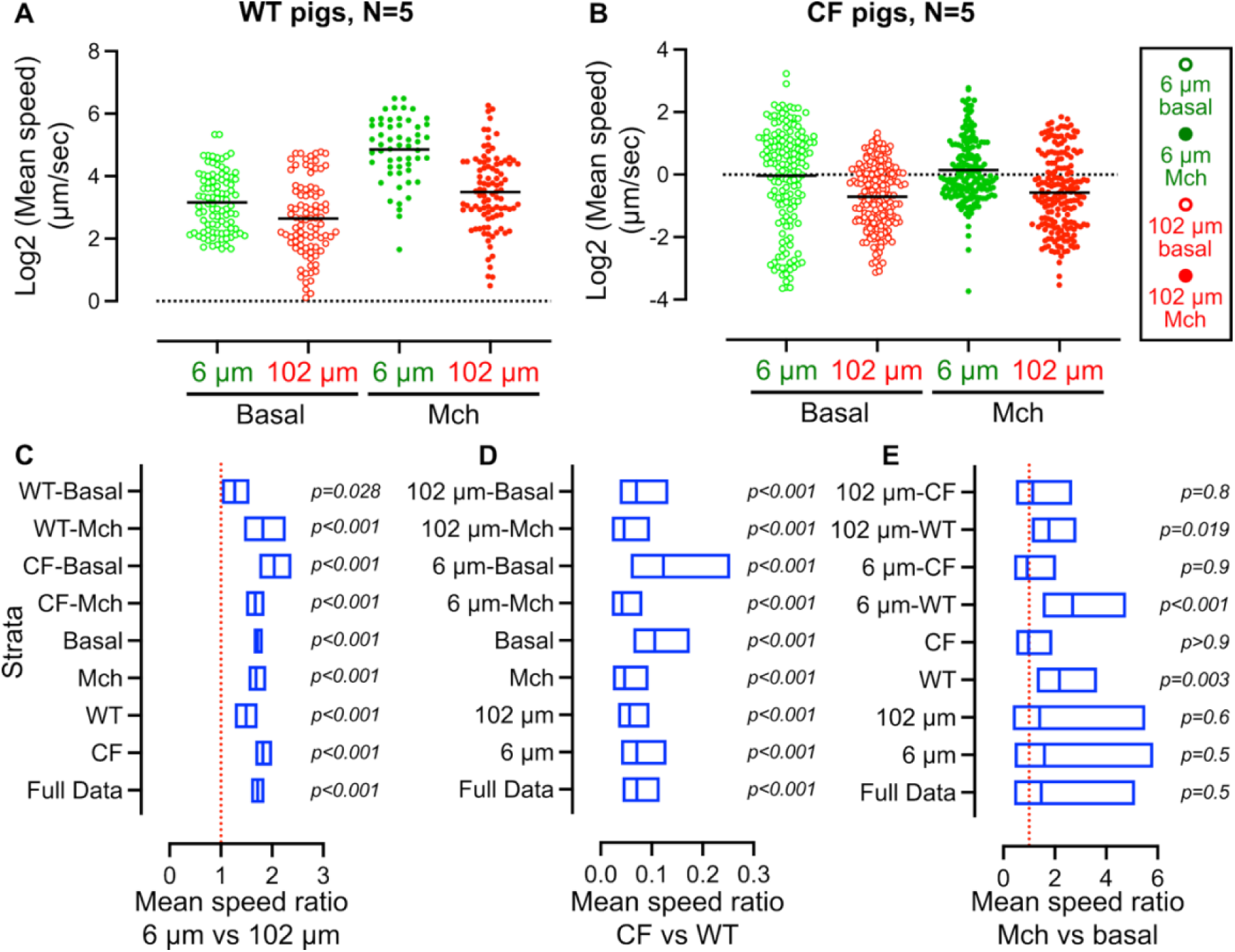
Particle size, CF airway disease, and Mch stimulation impact the mean speed of particle motion on the airways. (A-B) The mean speed of microparticle motion on WT and CF airways. (C-E) Effect of particle size, CF airway disease, and methacholine stimulation on mean speed ratio. In panels A-B, each data point represents the mean speed of 30-50 particles over 1 minute. In panel C-E, data are represented by mean speed ratio, 95% confidence intervals, and p-values. Generalized linear mixed models were used to assess the effects of genotype (CF vs WT), particle size (6 µm vs 102 µm), and MCH (Mch vs basal) on particle mean speed. N=5 pigs for each condition.

On CF airways, both 6 µm and 102 µm particles had significantly reduced mean speed compared to WT airways (Fig. 4B, note the difference in y-axis from Fig. 4A), yielding a mean speed ratio <1 for all strata (all p<0.001) (Fig. 4D). The impaired motion of 102 µm particles was consistent with previous *in vivo* MCT studies using tantalum micro-disks in CF pigs [13]. Here, we demonstrate that 6 µm particles, similar in size to inhaled bacteria, exhibit reduced clearance and speed. Furthermore, although Mch increased the mean speed of particles on WT airways, it did not show a significant effect on CF airways (Fig. 4E). Alongside the mean speed, the maximum (Fig. S2), median (Fig. S3), and minimum speed (Fig. S4) of particles were compared, which revealed the same trend as particle clearance and mean speed results (Table S1).

### 6 µm and 102 µm microparticles have distinct motion, causing differences in MCT

We sought to explain the difference in MCT of 6 µm and 102 µm particles by correlating their motions with the mucus structures that enable their movement. Recent studies revealed heterogeneous structures and motions of airway mucus. On pig airways, Ostedgaard *et al.* found that mucin MUC5B forms strands that emerged from SMG ducts, MUC5AC forms threads that emerged from airway surface goblet cells, and MUC5AC threads often coat MUC5B strands [22]. Ermund *et al.* demonstrated that such bundles of mucus transport more slowly than the airway surface liquid flow [23]. Hoegger *et al.* revealed that mucus strands failed to break free from SMGs and accumulated on CF airway surfaces [4]. Thus, we analyzed how individual particles move with airway mucus.

Under WT basal conditions, 6 µm particles move instantaneously. Most of them aggregate into strand-like shapes, capture 102 µm particles, and propel them from the airway surface (Fig. 5A-B). Before being captured by adjacent strands, 102 µm particles remain static (yellow arrowhead, Fig. 5A-B), causing a lag in motion. Once captured, the speed of 102 µm particles is similar (yellow arrowhead, Fig. 5A) or less (yellow arrowhead, Fig. 5B) than that of 6 µm particles in strands. If some 102 µm particles are not captured, they move trivially (blue arrowheads, Fig. 5B). We quantitatively analyzed the speed of microparticles that were trapped in a strand-like shape or not under WT-basal conditions (Fig. 5C). We confirmed that the mean speed of microparticles trapped in a strand-like shape was larger than that of those not trapped, suggesting that the mucus strand enhanced the MCT of microparticles. Furthermore, 102 µm particles did not move faster than 6 µm particles under all circumstances. Under Mch conditions, we occasionally found that 6 µm and 102 µm particles moved without aggregating into a strand-like shape; instead, they moved in the same direction and speed (Fig. S5). The particle aggregation into strand-like shapes is consistent with findings on mucus structures [22] and supports the role of mucus strands in facilitating microparticle clearance.

**Fig. 5.**
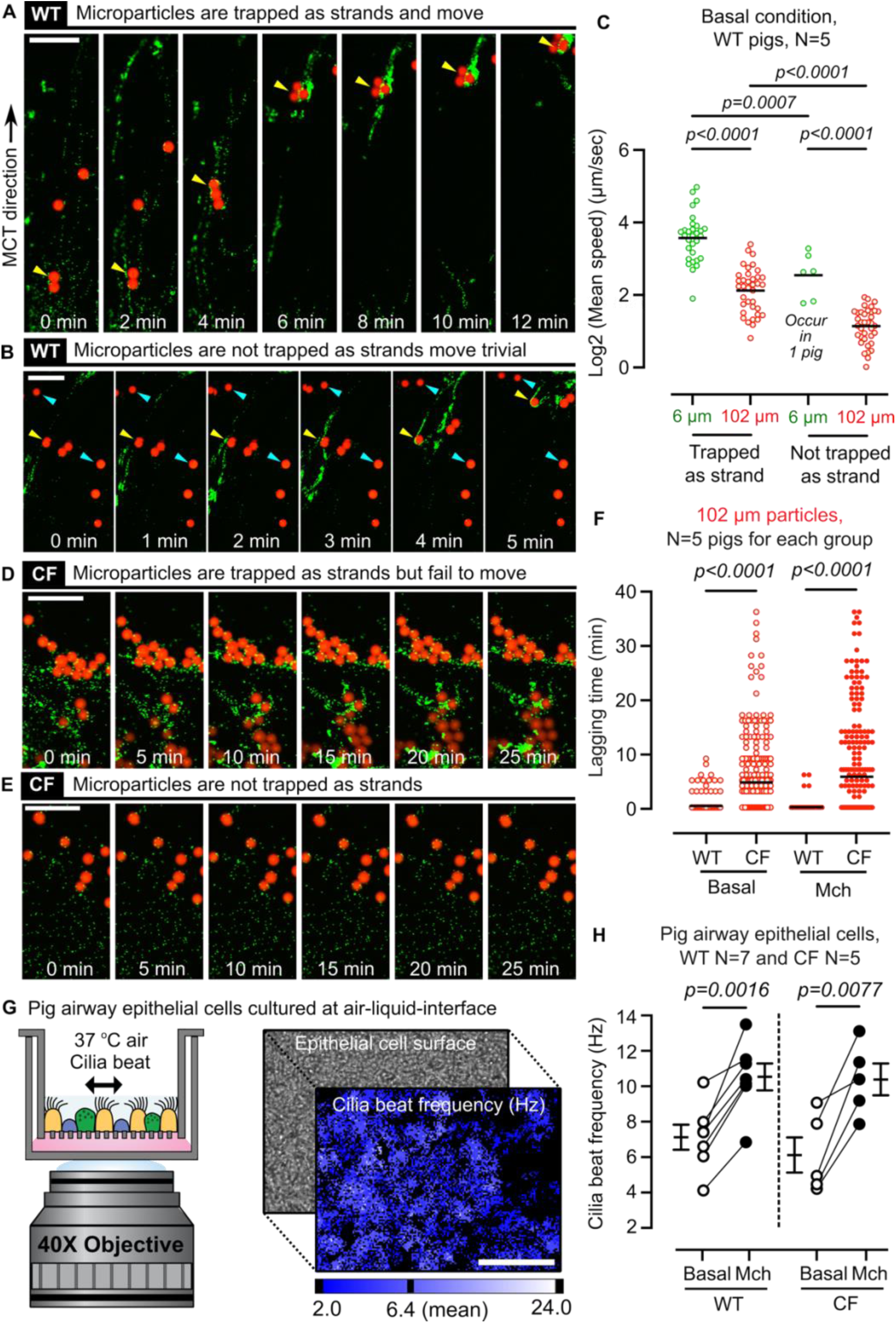
Motion of microparticles depends on their size, CF disease, and Mch stimulation. (A) Representative particle motion on WT airways, where 6 µm microparticles form strand-like shapes, trap 102 µm particles (yellow arrowhead), and clear them from the airway surface. (B) Representative particle motion on WT airways, where 102 µm particles that are trapped in strands of 6 µm microparticles and move (yellow arrowheads); and 102 µm microparticles move trivially without being trapped as a strand (blue arrowheads). (C) Mean speed of microparticles on WT airways with and without being trapped as strands. (D) Representative particle motion on CF airways, where 6 µm particles form as strand-like shapes, and 102 µm particles aggregate around the strands, but both fail to clear. (E) Representative particle motion on CF airways, where 6 µm and 102 µm particles move trivially, and no strands form. (F) Lagging time before the motion of 102 μm particles starts on WT and CF airways. (G) Method of studying cilia beat frequency on primary airway epithelial cell cultures. (H) Cilia beat frequency on WT and CF epithelia with/without basolateral Mch addition. In panels A, B, D, and E, the scalebars are 500 µm. In panel G, the scalebar is 200 µm. In panel C, each data point represents the mean speed of ∼50 particles over 1 minute. In panels C and F, One-way ANOVAs are used for statistical comparison, and N=5 pigs for each group. In panel H, paired T-test was used for statistical comparison, N=7 for cultures of WT pig epithelial cells, and N=5 for cultures of CF pig epithelial cells.

To further illustrate the size effect, we studied the MCT of tantalum balls (∼500 μm in diameter) and polystyrene nanoparticles (20 nm in diameter) on trachea explants submerged in Krebs buffer (N=3 WT pigs). The use of metallic balls and nanoparticles exaggerated the size differences, and liquid submersion allowed better observation of mucin structures. We found that nanoparticles quickly flowed across the airway (Movie S2). Most nanoparticles flowed individually or as small clusters, suggesting that they were carried by airway surface liquid or mucus threads. Some nanoparticles bound to mucus strands, accumulated around the metal ball, and initiated the movement of the metal ball after a lag time. The speed of the metal ball was significantly slower than that of nanoparticles in free liquid; it was also similar to or slower than that of nanoparticles that attached to mucus strands. The motion difference between large and small particles on submerged airways is consistent with our observations on airways with natural mucus layers.

### Microparticles had slower motion and prolonged lagging, impairing MCT on CF airways

On CF airways, many 6 μm particles aggregated as a strand-like shape, but failed to sweep 102 μm particles, causing particle aggregation on the airways (Fig. 5D). We also found that numerous particles move trivially and do not form strands (Fig. 5E). This observation explains the reduced particle speeds compared to WT airways. Besides reduced speeds, we also noticed that particles lagged before their motion started. The lagging time of microparticles was defined as the duration that a particle remained static before its motion began. Using 102 µm particles as an example, on WT airways, the 6 µm microparticles accumulate in a strand-like shape around the 102 µm particles, trapping and propelling them to start MCT. This process is quick on WT airways, resulting in a lagging time of ∼1 min before 102 µm particles started moving (Fig. 5F). However, due to the failed formation or motion of strands on CF airways, the lagging time of 102 μm particles (∼5 min on average) significantly increased. In addition, Mch did not appear to alter the lagging of 102 μm particles in CF airways.

### Methacholine increased cilia beating, influencing the MCT of microparticles

To explain the effect of Mch, we studied ciliary beating, which provides the driving force for MCT. We measured the frequency of ciliary beat using pig airway epithelial cells cultured at air-liquid interfaces and analyzed the data using Sisson-Ammons Video Analysis software (Fig. 5G). We found that Mch significantly increased cilia beat frequency on both WT (p=0.0016) and CF airways (p=0.0077) (Fig. 5H). Interestingly, the Mch increased particle clearance and speed only on WT airways, but not CF airways (Fig. 3 and 4). Previous studies also support these findings, where Mch facilitated mucus clearance on WT airways, but Mch caused mucus accumulation on submerged CF airways [4].

## Discussion

With the trachea-on-a-chip method, we revealed that MCT of microparticles is size-dependent, where 102 µm particles were cleared more slowly than 6 µm particles; MCT of microparticles was impaired on CF airways; and Mch increased MCT of microparticles on WT airways, but not CF airways. Our findings reveal a size-dependent particle clearance on normal and CF airways. Airways in humans and large mammals feature a thin layer of airway surface liquid with mucin MUC5AC threads from goblet cells, and MUC5B strands primarily from SMGs [22]. We propose that small particles (*i.e.*, 6 μm) can be moved by either airway surface liquid, mucus threads, or mucus strands, whereas large particles (*i.e.*, 102 μm) require mucus strands accumulation (Fig. 6A). As a result, small particles move instantaneously and faster, large particles move slowly and lag before motion starts (Fig. 4 and 5F). Furthermore, mucus strands increase the motion of both large and small particles (Fig. 5C). The proposed mechanism explains previous observations, where tris-(2-carboxyethyl) phosphine fragments mucus strands, reducing the motion of micro-disks (∼ 500 μm) on pig airways [17]. We also proposed that reduced particle clearance in CF is due to SMG secretion of abnormal mucus [24] and airway surface liquid [25, 26], which produces a dense mucus layer (Fig. 6B). Small particles trapped in the mucus layer move more slowly, and large particles are captured in mucus strands that often fail to break free (Fig. 6C) [4, 24]. The effect of Mch suggests a collaborative role between cilia beating and mucus in determining MCT (Fig. 6C). We speculate that, on normal airways, increased cilia beating accelerates the motion of airway surface liquid and mucus threads that carry small particles; it also increases detachment of mucus strands that carry larger particles.

**Fig. 6.**
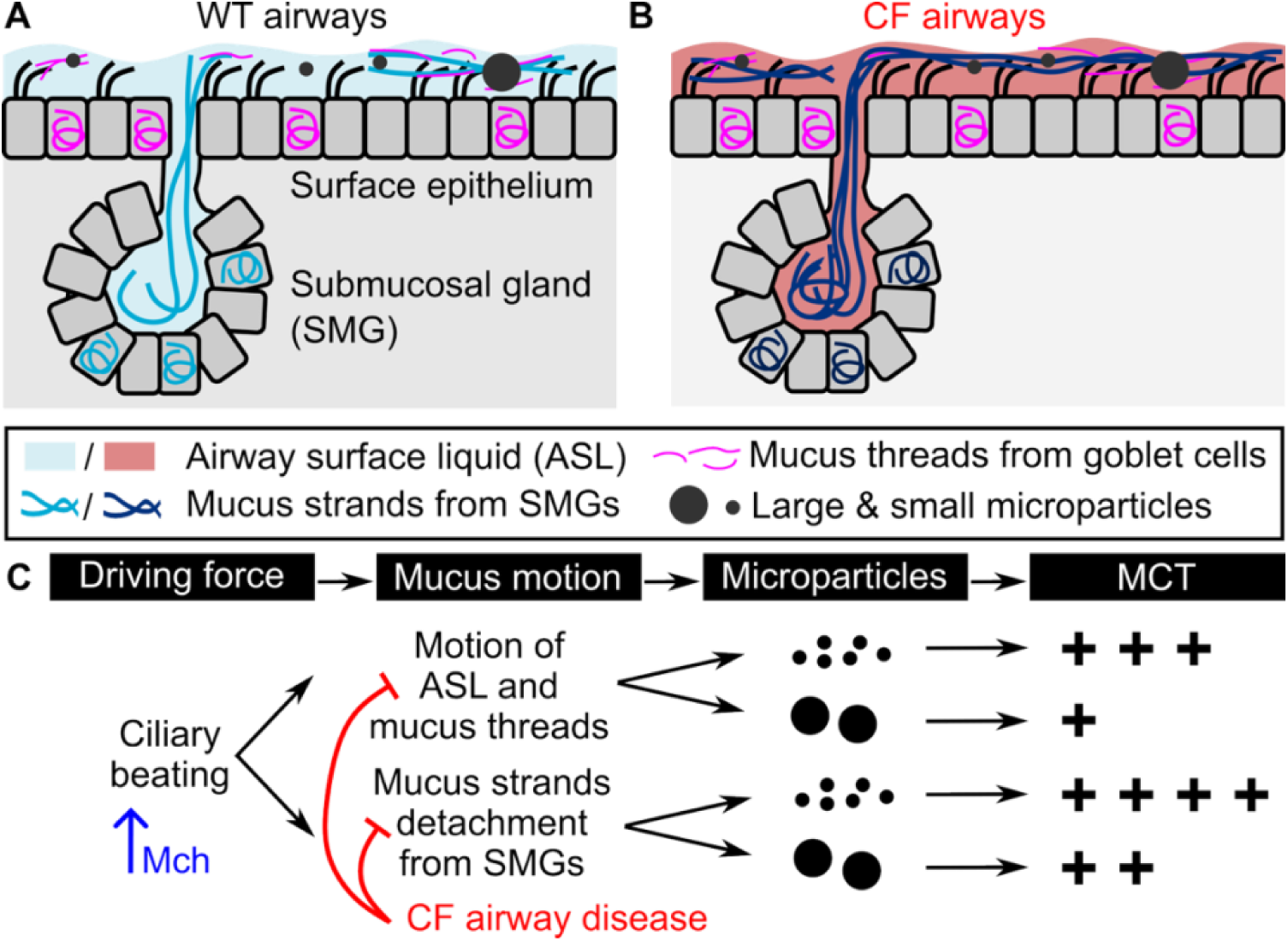
A model of size-dependent particle MCT on airways. (A) On WT airways, small particles carried by airway surface liquid and mucus threads move faster, and small particles carried by mucus strands move slower. Large particles are cleared through mucus strand breakage, resulting in slower movement and lagging time for mucus strand tethering. (B) On CF airways, abnormal airway surface liquid and failed breakage of mucus strands from submucosal gland duct significantly reduce MCT of microparticles. (C) Schematic demonstration of the effect of Mch and CF on MCT of microparticles with different sizes.

When increased cilia beating is powerful enough to move large particles directly, it causes a non-strand-based particle motion on WT airways (Fig. S5). However, CF mucus with abnormal elasticity and tensile strength [4, 24] fail to detach even under increased cilia beating, where Mch became ineffective.

Our results may have implications for airway disease pathogenesis, diagnostics, and therapeutics. The requirement of mucus strands in clearing large particles suggests the importance of SMGs in airway defense. In CF airways, we showed that Mch, a cholinergic agonist [24], elicits cilia beating but fails to enhance particle clearance. The accumulation of mucus and the presence of trapped bacteria may together cause infection and initiate the vicious cycle of CF lung disease.

Therapeutically, since MCT can clear drug-carrier particles, our results suggest that to achieve specific drug efficacy for therapeutic purposes, the size and other properties (*e.g.*, shape and surface properties) of the carrier need to be carefully evaluated. Our results also help to interpret diagnostic assays where various tracer particles have been applied *in vivo* [7]. Previous works have reported a wide range of data regarding particle residence times in the airways, and they attributed the disparity to variations in particle deposition positions [1, 2, 7]. In this work, we normalized the particle deposition and showed that particle motion is size-dependent. Based on our findings, we speculate that distinct particle clearance results hold; however, simple comparison across reports may not be valid.

The trachea-on-a-chip method is advantageous in studying the MCT of microparticles. First, apical perfusion of humidified O_2_/CO_2_ maintains a natural mucus layer for MCT, and the basolateral perfusion maintains normal tissue function, eliminates metabolic waste, and stimulates mucus production [19]. Second, the trachea-chip method features superior spatial (∼5 μm) and temporal (∼1 min) resolution to track the motions of 6 μm particles individually. In contrast, previous methods applied large Teflon [12] or tantalum disks [13] to the airways. Our findings showed that the motions of large and small particles are different. Third, previous lung/airway-on-a-chip methods successfully integrate the epithelial, endothelial, and immune cells [27, 28] with micro-fabricated device to study the airway defense function [29] and CF disease [30], here we integrated tissue explants from newborn pigs to study airway particle clearance. Pigs have similar trachea anatomy and cellular composition to humans [18], airway explants preserve organ geometry, anatomic architecture (*e.g.*, SMGs), and cell types. Newborn CF pigs avoid secondary disease manifestations from infection or inflammation, thereby identifying defective MCT at birth. Other genetically or environmentally modified pig models can be tested in the future, such as ectodysplasin gene knock-out pigs, which lack SMGs throughout the airways [31].

This work inevitably has limitations that can be addressed in the future. We tested 6 µm and 102 µm particles that mimic inhaled bacteria and radioactive tracers, respectively. Future studies are expected to answer how other properties of particles impact MCT. For example, particles and certain bacteria [32] have hydrophilic polymer coatings (*e.g.*, polyethylene glycol) that exhibit minimal adhesion to porcine mucus [33], allowing these particles to penetrate the mucus layer [34–36]. Second, this work tested the effect of Mch; future studies would study MCT under various airway physiological states. For example, tuning flow rates of the basolateral flow to generate airway deformation that mimics breathing and coughing. Since many ion channels (*e.g.,* CFTR) are mechano-sensitive [37–39], we expect that mechanical perturbations will impact MCT. The device can also adjust the apical H_2_O saturation level to study MCT under dehydrated conditions [40]. Third, future studies could benefit from integrating human tissues. While tissues from newborn pigs provide insight into the early pathogenesis of airway disease, we expect that tissues from human donors would yield knowledge on advanced or end-stage diseases.

In summary, we demonstrated a particle size-dependent MCT mechanism on normal and CF airways with a trachea-on-a-chip device. The fast, instantaneous motion of small particles and the slow, lagging motion of large particles suggest they were cleared by distinct MCT mechanisms. We anticipate that the trachea-chip method and findings on MCT of microparticles will have a broad impact on future research in airway disease pathogenesis, diagnostics, and therapeutics.

## Methods

### Device fabrication

The trachea-on-a-chip device was designed using Creo 10.0 software (PTC, USA) and fabricated via computer numerical control (CNC) machining (Protolabs, USA) from 7075 aluminum.

### Reagents

A HCO ^-^/CO_2_ buffered Krebs-Ringer saline containing (in mM): 118.9 NaCl, 25 NaHCO_3_, 10 dextrose, 2.4 K_2_HPO_4_, 0.6 KH_2_PO_4_, 1.2 CaCl_2_, 1.2 MgCl_2_, pH=7.4 with 5% CO_2_ was used. Mucus secretion and cilia beating were stimulated with basolateral perfusion of methacholine (1.28×10^-5^ mol/L, Sigma-Aldrich, USA). Fluorescent dry particles with diameters of 6 μm (Fluoro-Max Green Fluorescent Polymer Microspheres, ThermoFisher Scientific, USA) and 102 μm (Fluoro-Max Red Fluorescent Polymer Microspheres, ThermoFisher Scientific, USA) were used to study their clearance *via* MCT.

### Animals

Cystic fibrosis transmembrane conductance regulator knock-out (*CFTR^-/-^*) pigs (*i.e.*, CF pigs) were generated previously [41], and obtained from Exemplar Genetics. Wildtype (*i.e.*, WT) pigs were purchased from either Exemplar Genetics or Premier Biosource. Both WT and CF pigs were studied less than 3 days after birth. Pigs with both genders are used in this work, because sex was not considered as a biological variable in particle clearance. Animal anesthesia was administered using ketamine (20 mg/kg, I.M.) (Phoenix Pharmaceutical, Inc., USA) and xylazine (2 mg/kg, I.M.) (Lloyd, USA). Euthanasia was achieved with I.V. Euthasol (Virbac, France). A tracheal segment between the opening of the right cranial lobe and the larynx was cut with a surgical blade between cartilage rings. After being removed from pigs, the tracheal segments were immediately placed on surgical gauze saturated in Krebs-Ringer buffer and stored at 4 °C for less than 24 hours before experimentation. Notably, the tracheal tissue was never submerged in excessive liquid.

### Application of microparticles

Microparticles with diameters of 6 μm (green fluorescent) and 102 μm (red fluorescent) were applied to the trachea surface *via* a homemade nebulizer (Fig. S1). The nebulizer was constructed with a 1 mL pipette tip inserted into an oval blood pressure bulb. A second 1 mL pipette tip was subsequently placed firmly onto the pipette-bulb apparatus for particle loading. After the particles were loaded into the secondary pipette tip, they were ejected from the pipette onto the tracheal surface by squeezing the bulb. In experiments characterizing the density of particle application, we sandwiched a piece of 3M tape, instead of a trachea explant, into the trachea-on-a-chip device. The surface of the tape adheres to particles and lacks MCT, thereby preventing particle motion.

### Visualization of MCT of microparticles

After application of microparticles, the apical chamber was sealed with a cover piece made of polymethyl methacrylate. Air with 100% humidity and 5% CO_2_ was perfused apically, and Krebs-Ringer buffer was perfused basolaterally with a syringe pump (flow rate 100 µL/min) (NEMESYS, CETONI GmbH, Germany) until the end of the experiment. In some experiments, 2.5 µg/mL methacholine (Sigma-Aldrich, Japan) was perfused basolaterally to stimulate mucus secretion and cilia beating. The entire device was placed in an environmental chamber at 37 °C and then mounted in an upright confocal microscope (Nikon A1R, USA). Motion of fluorescent microparticles was visualized using a 4X (NA=0.13) dry objective lens (Nikon Plan Fluor, Japan). Excitation/emission wavelength to visualize 6 μm green particles were 468/508 nm, and 102 μm red particles were 542/612nm. To assess particle motion across the entire tracheal surface in the chamber, the confocal microscope was synchronized to a stage controller (MS-2000-500, Applied Scientific Instrumentation, USA). The stage controller moved the microscope stage to different positions on the tracheal surface to take panoramic images. In this study, 4 fields along the ventral-dorsal direction and 6 fields along the proximal-distal direction were stitched together for a single panoramic image that contains 24 individual images. At each imaging field, bright field images (to visualize the tracheal surface), red-fluorescent images (to visualize 102 μm particles), and green-fluorescent images (to visualize 6 μm particles) were taken sequentially. The panoramic images were recorded every minute for a total of 36 minutes.

### Calculation of microparticle clearance

The time-lapse of panoramic images was processed using Imaris 10 software (Oxford Instruments, Switzerland) to track particle motions. Below, we exemplify the protocol using 102 µm particles, where a similar protocol was employed for 6 µm particles. A region of interest (ROI, 2 mm by 2 mm) was selected on the airway surface. The ROI was selected on the apical chamber without the edges, and the ROI remained unchanged during the 36 minutes for particle analysis. To track particle motion, 102 µm microparticles were first identified using the Imaris software based on their size, and a manual double-check was applied to add or remove any faulty recognitions. After 102 µm particles were recognized throughout all frames, particles were linked every 1-minute interval to obtain the motion trajectories. A manual double-check was applied to remove trajectories that did not link correctly.

To calculate the clearance of 102 µm particles, the number of particles at T_0_ was counted. The particle trajectories revealed the time at which the particle left the ROI, thus allowing the computation of the number of particles that were initially in the ROI and then left the ROI at certain time points. The percent of particles that leave the ROI between T_0_ and T_1_ relative to the number of particles in the ROI at T_0_. When calculating the particle clearance, we only quantified particles that were originally in the ROI and ignored particles that entered the ROI after T_0_. We calculated particle clearance each minute and evaluated 3 independent factors (*i.e.*, 6 µm vs. 102 µm, basal vs. Mch, and WT vs. CF), resulting in 8 experimental groups. For each group, results from 5 pigs were plotted vs time (Fig. 3A-B).

### Calculation of microparticle speed

To calculate the speed of particles, the trajectories of particles that interfere (i.e., enter, remain, and leave) with the ROI (2 mm by 2 mm) were analyzed. At an arbitrary time between T_i_ and T_i+1_, speeds of particles (*v_i_*) were obtained from their trajectory by the displacement of each particle divided by the duration of 1 min. Between T_i_ and T_i+1_, *v_i_* forms a set

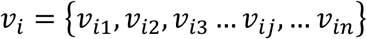

where *n* is the number of particles whose speed was analyzed. In general, 30-50 microparticles were counted. The maximum, minimum, and median speeds of *v_i_* were collected using Imaris software.

And the mean speed of particles (*v̅*_*l*_) was calculated by

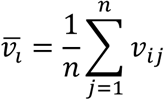

In a typical particle tracking process with a duration of 36 minutes, the mean speed at each 1-minute time point further forms a set (*v_mean_*) that contains 36 elements. For some WT airways, microparticles were cleared within 10 min, thus the *v_mean_* contains elements number less than 36.

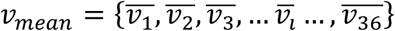

The *v_mean_* was calculated at each minute with 3 independent factors (*i.e.*, 6 µm vs. 102 µm, Mch vs. basal, and CF vs. WT), resulting in 8 experimental groups. For each group, results from 5 pigs were plotted (Fig. 4A-B). Similarly, the maximum, minimum, and median speed from 5 pigs were plotted (Fig. S2-S4).

### Calculation of microparticle speed with and without strands

To study how the aggregation of particles into strands impacts their motion, we first observed the “strand-like shape” using 6 µm microparticles, where a group of microparticles aggregated into a continuous, elongated structure that moved along the airway surface. Similar strands were observed previously using 20 nm nanoparticles on submerged airway surfaces [4]. To calculate the speed of 6 µm microparticles that move with strands, we tracked microparticles that were located at the middle of the strand using Imaris software. To calculate the speed of 102 µm microparticles that move with strands, we tracked a particle that was surrounded by 6 µm microparticles with a strand-like shape. The speeds of particles with and without a strand were analyzed in an ROI of 2 mm by 2 mm on the airway surface. On each airway, 4 ROIs were chosen to calculate the speed of microparticles. The speed of microparticles was analyzed every minute until 36 minutes or until they were cleared from the ROI. In each 1 minute, the mean speed was calculated as described in the previous section. We reported the log2(mean speeds) of microparticles on 5 WT pig airways under basal conditions (Fig. 5C).

### Calculation of lagging time before microparticle motion

The lagging time of 102 µm microparticles was computed with Imaris software. To determine the states of station and motion, particle trajectories data generated with Imaris were analyzed. The static condition was defined if a particle did not change its position over time, and a particle in motion was defined if a change in particle position was detected. The time point from static to motion was used as the lagging time. We analyzed the lagging time in ROI of 2 mm by 2 mm on WT and CF airways and reported (Fig. 5F).

### Motion of Tantalum balls and nanoparticles on submerged airways

A segment of pig trachea (length of ∼1 cm) was dissected from newborn pigs, opened along the ventral surface, flattened to expose the mucosal surface, and pinned with dental wax in a 35 mm circular petri dish. The trachea explant was submerged in 40 mL of Krebs buffer at 37 °C. Methacholine (100 µM) was added to the solution to increase the production of mucus strands. 2-4 µL nanospheres (20 nm, red fluorescence) were added to label strands of mucus. Metallic spheres (Tantalum, 500 µm diameter, Bal-Tec, USA) were manually added to the trachea surface using a P200 pipette tip. A confocal microscope with a 10X objective was used, where a reflected channel was used to visualize the movement of the metallic sphere, and a red fluorescent channel was used to visualize the movement of nanospheres and mucus strands. We conduct experiment on 3 WT pig airways and reported the movies of particle motions (Movie S2).

### Cilia beat frequency

Primary cultures of airway epithelia were derived from cells harvested from WT and CF pig ethmoid tissues. Epithelial cells were enzymatically isolated, seeded onto collagen-coated inserts (Costar 3470, Corning, USA), and cultured at the air-liquid interface. The differentiated epithelia were used for ciliary beat frequency assays after a minimum of 14 days had passed since seeding. Cell culture medium comprised a 1:1 mixture of Dulbecco’s modified Eagle medium/F-12, supplemented with 2% Ultroser G, 5 µg/mL insulin, 0.5 ng/mL epidermal growth factor, 0.125 µM transferrin, 0.5 mg/mL albumin, and 50 nM retinoic acid. One day prior to the ciliary beat frequency measurement, we used PBS (with Ca^2+^ and Mg^2+^) to wash the apical side of the air-liquid interface culture. During the assay, air-liquid interface culture was put on top of the 35 mm glass-bottom dish at 37°C. 1 mL of culture media, with 0.1% PBS or 2.5 µg/mL methacholine, was added to the basolateral side. Videos of ciliary motion were obtained using the microscope (Zeiss Axio Observer, Germany) with a 40X water immersion objective (NA=1.2) at 50 frames per second. Each video contains at least 128 frames (duration ∼2.6 sec), and a minimum of eight videos were recorded in random positions of each culture. The ciliary beat frequency (CBF) was analyzed using video analysis software (SAVA; Ammons Engineering, Mt. Morris, MI, USA), where a mean value was calculated from the field of the video. We reported results of cilia beat frequency measurement on 7 WT pigs and 5 CF pigs (Fig. 5H).

### Statistics

To quantify the effect of genotype, particle size, and Mch on particle clearance, Prism 10, R v4.3.1, and SAS v9.4 were used. Cox proportional hazards models were used to assess the effects of genotype (WT vs. CF), particle size (6 µm vs 102 µm), and methacholine (basal vs. Mch) on time of particle clearance on the airways. Particles that did not clear within 36 minutes were censored at that time. In all models, a random effect was included to account for inherent within-subject correlation between particle measurements from the same pig. Assessments between factor levels were made on the full data set as well as on datasets restricted to the other factors. All modeling results are reported via clearance ratio (*i.e.*, Hazard Ratios in Cox proportional hazards models), 95% Confidence Intervals, and P-Values (Fig. 3C-E). A p-value less than 0.05 was considered statistically significant.

To quantify the effect of genotype, particle size, and Mch on particle speed, Prism 10, R v4.3.1, and SAS v9.4 were used. Generalized linear mixed models were used to assess the effects of genotype (WT vs. CF), particle size (6 µm vs 102 µm), and methacholine (basal vs. Mch) on particle speed. Separate models were constructed for the minimum, maximum, mean, and median particle speeds during each 1-min time interval of measurements. Due to the right-skewed nature of these outcomes, Gamma models with a log link were used. A shift of 0.001 was applied to outcomes to ensure all values were greater than 0. In all models, a random effect was included to account for inherent within-subject correlation between repeated measurements from the same pig. Assessments between factor levels were made on the full data set as well as on datasets restricted to the other factors. All modeling results are reported via Mean Ratios, 95% Confidence Intervals, and P-Values (Fig. 4C-E). A p-value less than 0.05 was considered statistically significant.

### Study approval

All animal studies were approved (#3071121) by the University of Iowa Animal Care and Use Committee (IACUC).

### Data availability

All data in the article are included in the Supporting Data Values file and supplemental materials. Request for additional information are addressed to the corresponding author.

## AUTHOR CONTRIBUTIONS

YX conceptualized the project; supervised the execution of the experiments; MS, KB, WD, MR, AF, YX carried out the experiments and performed the data acquisition; MS, YX drafted the manuscript; all authors read and approved the final manuscript.

## ACKNOWLEDGEMENTS

The authors would like to express their gratitude to Dr. Michael Welsh for his discussions on the project; University of Iowa tissue culture core and their staff members, including Phil Karp, Shu Wu, and Amy Clemons for providing access to tissue culture facilities; Animal core and their staff members, including Linda Powers, Nick Gansemer, Peter Taft, Mal Stroik for their assistance on cystic fibrosis pig tissue processing; Dr. Patrick Ten Eyck at University of Iowa Institute for Clinical and Translational Science for the statistical analysis of particle speed; and Tom Moninger for his help on microscopic observations.

