## Supplementary Info for "Particle size determines mucociliary transport mechanisms in normal and cystic fibrosis airways"

**SUPPLEMENTAL MATERIAL**

The supplemental material includes Figs. S1 to S5, and Movie S1 to S2 Caption. Other Supplementary Materials for this manuscript include Movie S1 to S2.

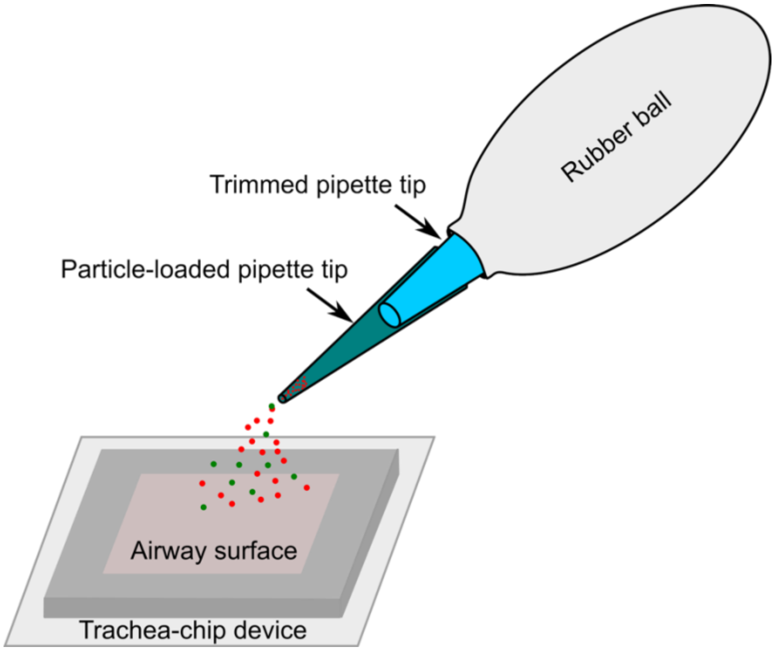


**Fig. S1. Device for particle application onto the airways.** A homemade particle application apparatus was used to spray dry microparticles onto the airway surface. It consisted of a rubber ball to apply pressure with hand, a trimmed 1 mL pipette tip to conduct air flow, and another disposable 1 mL pipette to backload the dry particles and spray them onto the airway surface.

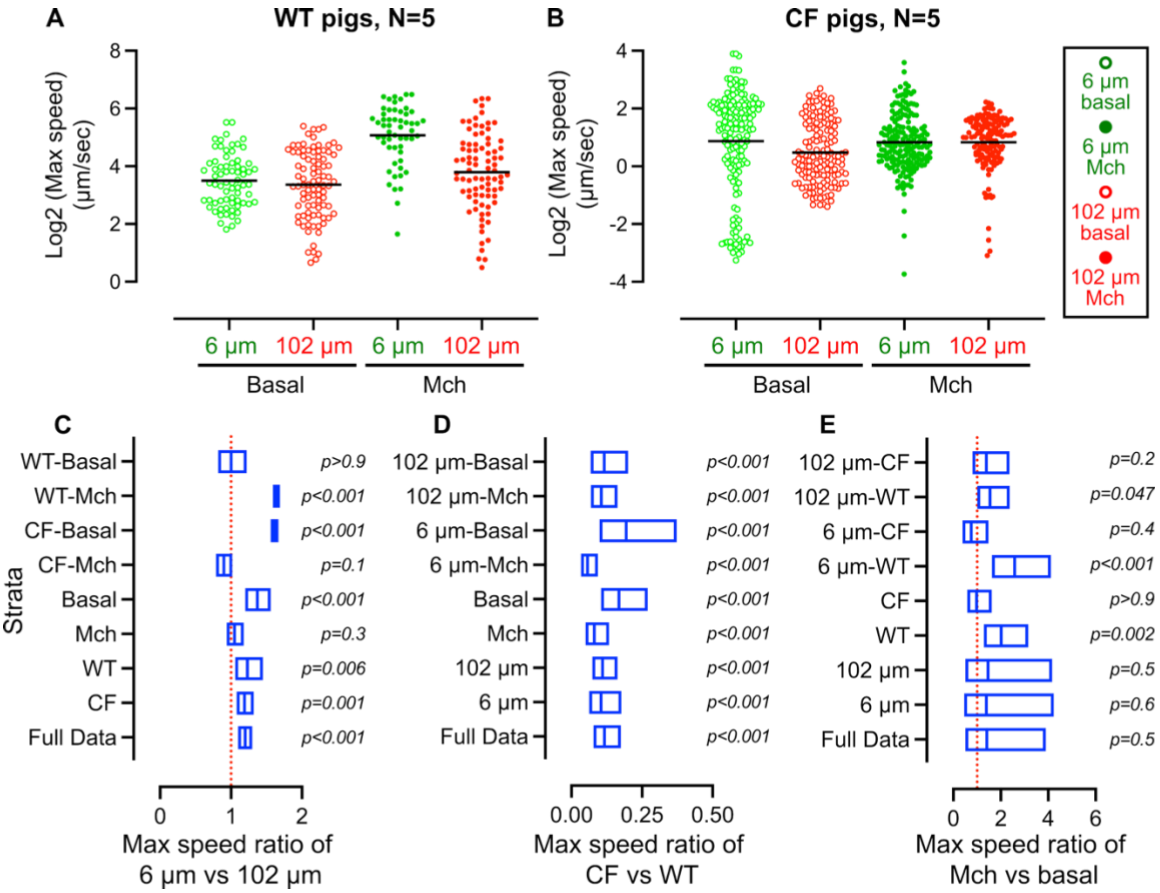


**Fig. S2. Particle size, CF airway disease, and Mch stimulation impact the maximum speed of particle motion on the airways. (A-B)** The maximum speed of microparticle motion on WT and CF airways. **(C-E)** Effect of particle size, CF airway disease, and methacholine stimulation on maximum speed ratio. In panels **A-B**, each data point represents the mean speed of 30-50 particles over 1 minute. In panel **C-E**, data are represented by max speed ratio, 95% confidence intervals, and p-values. Generalized linear mixed models were used to assess the effects of genotype (CF vs WT), particle size (6 µm vs 102 µm), and MCH (Mch vs basal) on particle max speed. N=5 pigs for each condition.

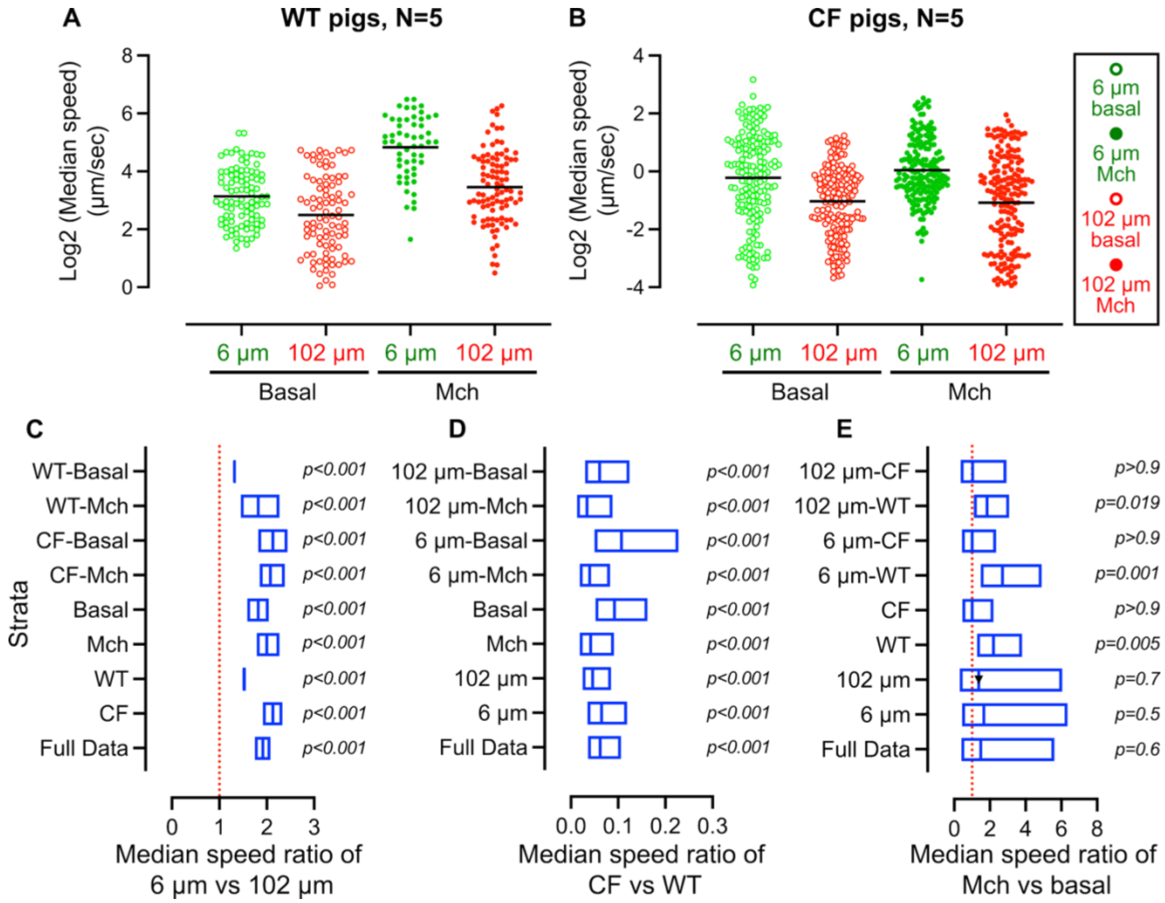


**Fig. S3. Particle size, CF airway disease, and Mch stimulation impact the median speed of particle motion on the airways. (A-B)** The median speed of microparticle motion on WT and CF airways. **(C-E)** Effect of particle size, CF airway disease, and methacholine stimulation on median speed ratio. In panels **A-B**, each data point represents the mean speed of 30-50 particles over 1 minute. In panel **C-E**, data are represented by median speed ratio, 95% confidence intervals, and p-values. Generalized linear mixed models were used to assess the effects of genotype (CF vs WT), particle size (6 µm vs 102 µm), and MCH (Mch vs basal) on particle median speed. N=5 pigs for each condition.

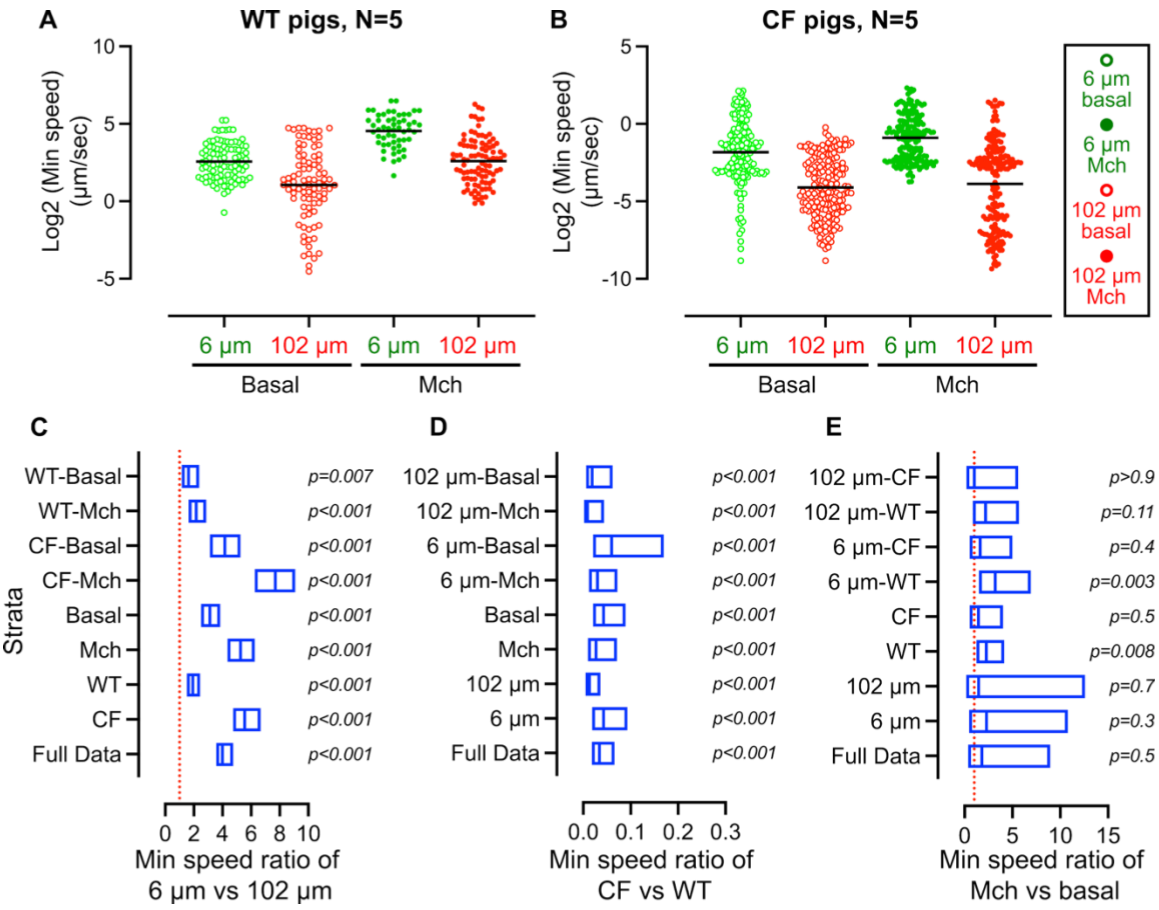


**Fig. S4. Particle size, CF airway disease, and Mch stimulation impact the minimum speed of particle motion on the airways. (A-B)** The minimum speed of microparticle motion on WT and CF airways. **(C-E)** Effect of particle size, CF airway disease, and methacholine stimulation on minimum speed ratio. In panels **A-B**, each data point represents the mean speed of 30-50 particles over 1 minute. In panel **C-E**, data are represented by minimum speed ratio, 95% confidence intervals, and p-values. Generalized linear mixed models were used to assess the effects of genotype (CF vs WT), particle size (6 µm vs 102 µm), and MCH (Mch vs basal) on particle minimum speed. N=5 pigs for each condition.

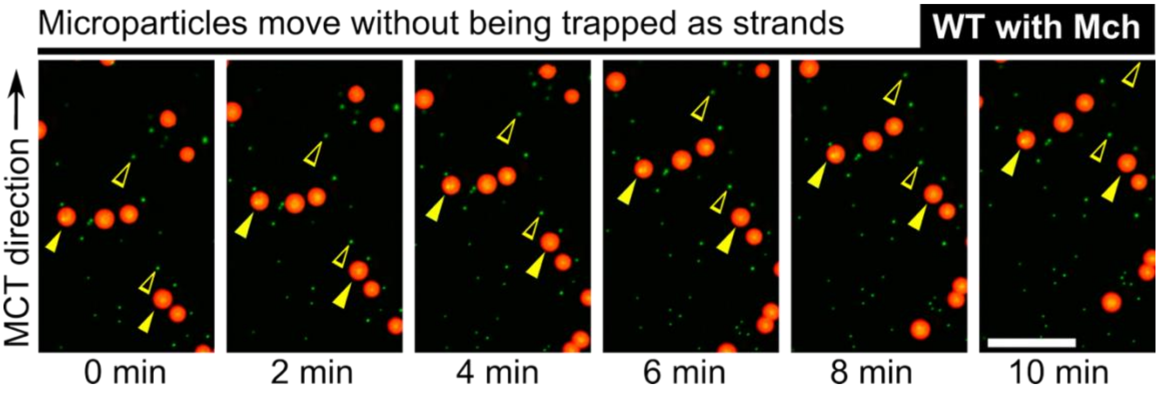


**Fig. S5. Representative image sequence of particle motion on WT airways with methacholine stimulation.** Upon Mch stimulation, both 102 µm (red) and 6 µm (green) microparticles move homogeneously without forming a strand shape.

**Table S1. P-value of particle clearance, mean, max, median, and min speed via MCT.**

| Comparisons | Groups | Particle clearance | Mean speed | Max speed | Median speed | Min speed |
|---|---|---|---|---|---|---|
| 6 vs. 100 μm | WT Basal | <0.001 | 0.028 | N.S. | <0.001 | 0.007 |
| | WT Mch | <0.001 | <0.001 | <0.001 | <0.001 | <0.001 |
| | CF Basal | <0.001 | <0.001 | <0.001 | <0.001 | <0.001 |
| | CF Mch | <0.001 | <0.001 | N.S. | <0.001 | <0.001 |
| Basal vs. Mch | WT 6 μm | <0.001 | <0.001 | <0.001 | 0.001 | 0.003 |
| | WT 100 μm | N.S. | 0.019 | 0.047 | 0.019 | N.S. |
| | CF 6 μm | N.S. | N.S. | N.S. | N.S. | N.S. |
| | CF 100 μm | N.S. | N.S. | N.S. | N.S. | N.S. |
| CF vs. WT | Basal 6 μm | <0.001 | <0.001 | <0.001 | <0.001 | <0.001 |
| | Basal 100 μm | <0.001 | <0.001 | <0.001 | <0.001 | <0.001 |
| | Mch 6 μm | <0.001 | <0.001 | <0.001 | <0.001 | <0.001 |
| | Mch 100 μm | <0.001 | <0.001 | <0.001 | <0.001 | <0.001 |

*Significant changes (P<0.05) are highlighted in blue.*

**Movie S1. Mucociliary transport of microparticles on WT and CF airway surfaces.** Clip A. Schematic of experimental setup. Clip B. MCT of fluorescent microparticles on a WT airway surface (duration is 36 min). Clip C. MCT of fluorescent microparticles on a CF airway surface (duration is 36 min).

**Movie S2. Mucociliary transport of a metal ball and nanoparticles on a submerged airway surface.** Clip A. Schematic of experimental setup. Clip B. MCT of a 500 µm tantalum ball and 20 nm particles on a submerged WT airways surface (N=3 WT airways).
